# First Record of *Aedes (Stegomyia) albopictus* (Diptera: Culicidae) in the state of Acre, Brazil

**DOI:** 10.1101/2022.08.18.504440

**Authors:** Ricardo da Costa Rocha, Acigelda da Silva Cardoso, Janis Lunier de Souza, Eliana da Silva Pereira, Marcio Fernandes de Amorim, Maria Socorro Martins de Souza, Cleomar de Lima Medeiros, Maria Francisca Mendes Monteiro, Dionatas Ulises de Oliveira Meneguetti, Marcia Bicudo de Paula, Andreia Fernandes Brilhante, Tamara Nunes Lima-Camara

## Abstract

**Introduction:** *Aedes (Stegomyia) albopictus* (Skuse, 1854) was reported in Brazil for the first time in 1986, and has displayed successful expansion throughout the Brazilian territory.

**Methods:** During a routine activity to control dengue conducted by the Division of Entomology of the Municipal Health Department in Rio Branco, adults and immatures of Culicidae were collected in a peri-urban area.

**Results:** The Culicidae forms identified indicated that they belonged to the species *Ae. albopictus*.

**Conclusions:** This is the first official record of the presence of *Ae. albopictus* in the state of Acre, confirming its current presence in all Brazilian states.

---

*Aedes (Stegomyia) albopictus* (Skuse, 1894) is also termed the Asian tiger mosquito, with the original distribution in East Asia and the islands of the western Pacific and Indian Oceans (Bonizzoni et al., 2013). Currently, it occurs in all the continents of the world, barring Antarctica^1^. In Brazil, *Ae. albopictus* was first detected in 1986, in the state of Rio de Janeiro, in a batch of larvae collected from abandoned tires in the Federal Rural University of Rio de Janeiro campus itself, in the city of Seropédica^2^. Besides, in 1986, *Ae. albopictus* was recorded for the first time in the states of Minas Gerais and São Paulo and, in the following year, this species became established in all the regions of the southeastern states of Brazil^3^.

In 2003, the *Ae. albopictus* distribution was updated in Brazil and the seven states cited here, namely, Acre, Amapá, Piauí and Tocantins (Northern) and Ceará, Roraima and Sergipe (Northeastern), had not yet received the official record of the species^3^. Nearly, twelve years later, other Brazilian states also recorded the presence of *Ae. albopictus*, and its absence was noted only in the states of Acre, Amapá and Sergipe^4^. Towards the end of 2014 and in 2019, the presence of this mosquito vector was reported in the states of Sergipe and Amapá, respectively, with Acre being the only state having no official record of the presence of this species^5^.

In this work, the presence of *Ae. albopictus* is reported for the first time in the state of Acre. This is the first official record of the finding of immature forms (larvae and pupae) and adult females of the *Ae. albopictus* in Acre state.

During the routine activities performed to control dengue, by the Division of Entomology of the Municipal Health Department in Rio Branco, on March 31, 2022, *Ae. albopictus* pupae were collected from a peri-urban area of the city of Rio Branco, in a place classified as a high-risk, non-residential property, for the proliferation of vectors (Strategic Point-SP; 10° 00’ 22.8” S; 67° 50’ 57.7” W). After confirming the identification of the *Ae. albopictus* pupae, the Division of the Entomology team conducted new entomological research from April 8 to 13, 2022, covering an approximately 8-km radius from the initial point of the primary collection of the *Ae. albopictus* pupae (Figure 1). The peri-urban area research presented dense vegetation cover, a very significant factor for the proliferation of this species.

**Figure 1.**
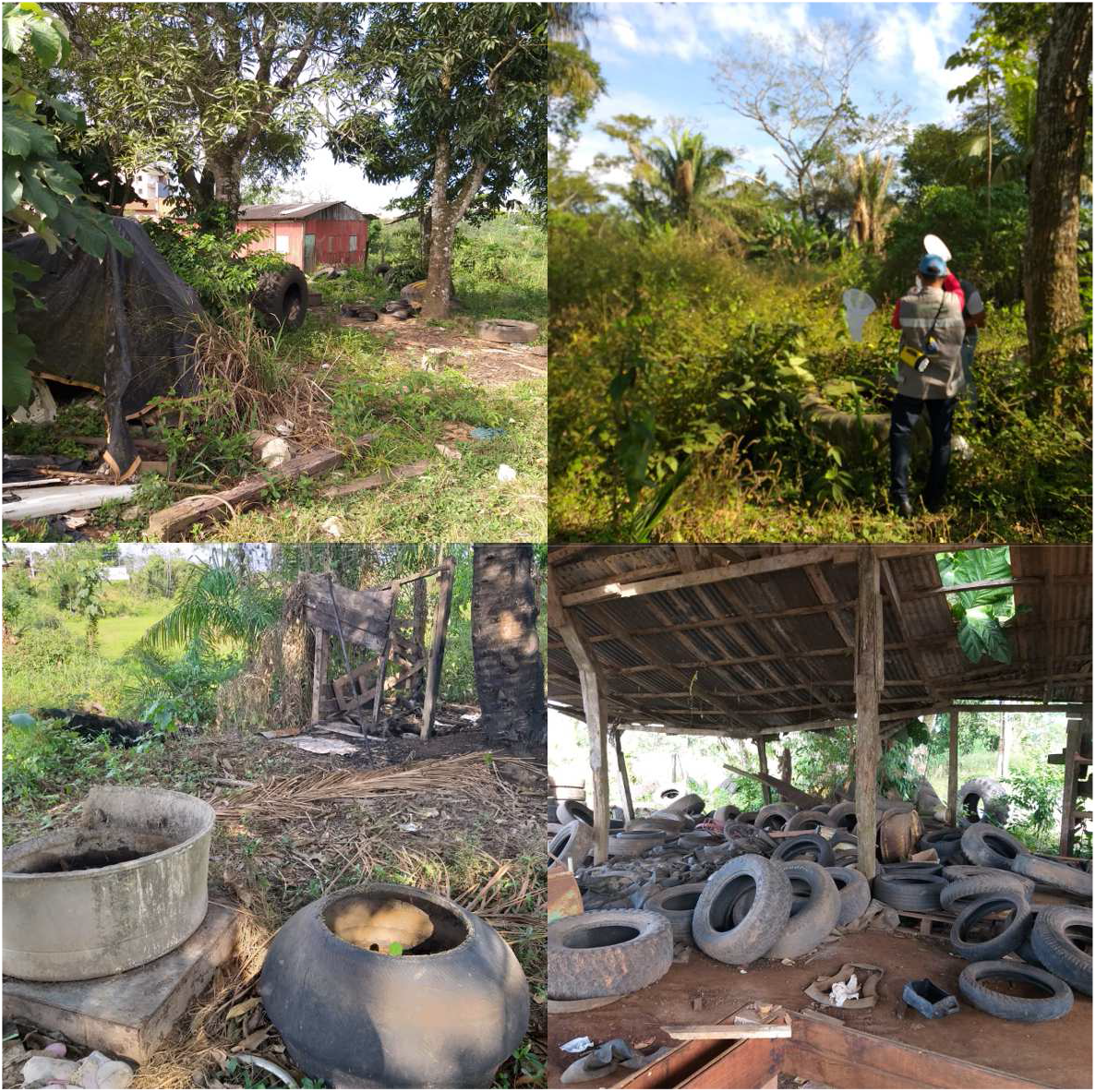
Collection area for immatures and adults of *Ae. albopictus*. Peridomiciliary area with high vegetation cover, but with the presence of garbage and discarded tires, demonstrating the presence of artificial breeding sites for the species. Rio Branco, Acre, Brazil.

In the second entomological survey, four female adults were collected, using a Pulsar-type net, and the larvae were collected with the help of a plastic pipette and stored in plastic tubes. The immature forms were gathered from different artificial breeding sites, such as tires and domestic waste, as well as from their natural breeding sites, like tree holes.

The specimens thus collected were sent to the Faculty of Public Health at USP, which confirmed that the species was *Ae. albopictus*, through morphological identification of the larvae, pupae and adult females. Photographs were taken using a Zeiss 2000-C Stereomicroscope and Zeiss AXIO Lab.A1 Microscope (Figures 2 and 3).

**Figure 2.**
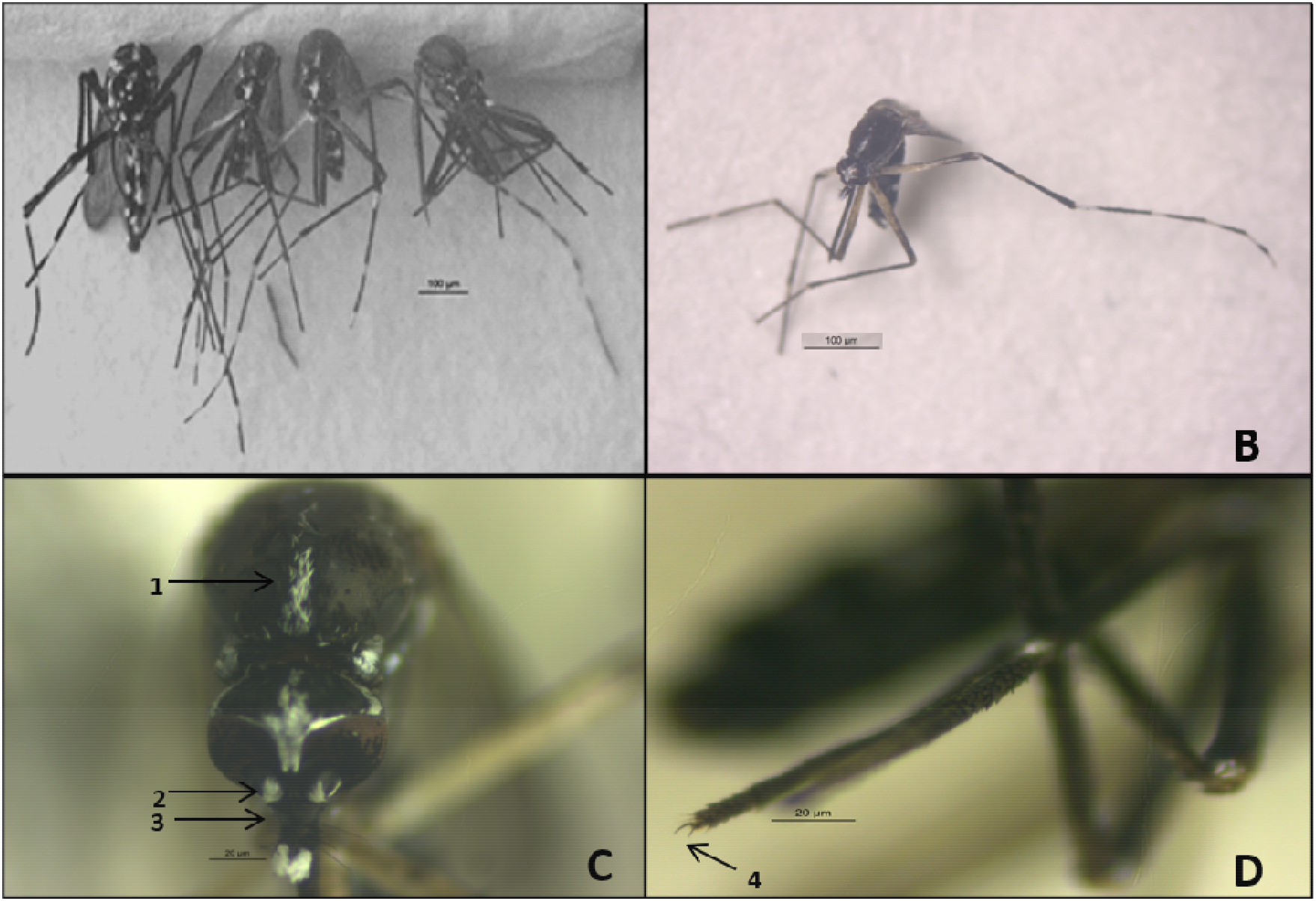
*Aedes albopictus*. A - Four females specimens. B - Dorsal view of the mesonotum. C - Head and Mesonotum: 1- longitudinal stripe of silvery scales; 2- silvery scales torus; 3- clypeus without scales. D - V tarsus of the anterior leg: 4- smooth shank nail.

**Figure 3.**
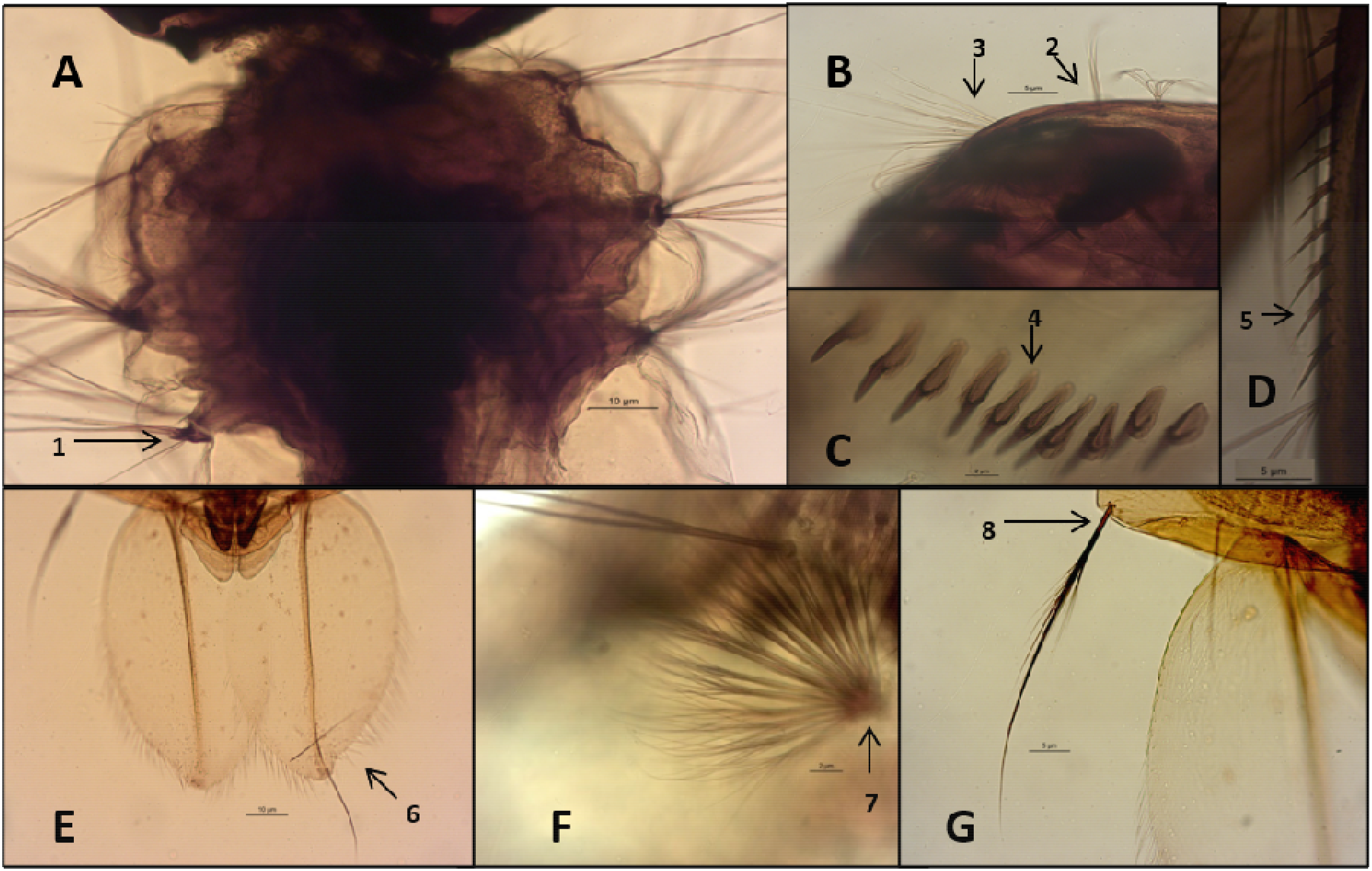
*Aedes albopictus*. A - Larvae thorax: 1- short lateral spines. B - Larvae head: 2- forked bristle 6; 3- bristle 7 multiple. C - VIII abdominal segment: 4- pecten in a single row, in the shape of a long spine and serrated base. D: 5- pecten siphon regularly spaced. E - Pupa 6- swimming reed with long hairs on the edge. F and G: Pupal abdomen: 7- bristle 1 with dichotomous bristles; 8- bristle 9 simple with small side hairs.

Figure 2 represents the four females examined, highlighting the longitudinal band of silvery scales in the mesonotum; on the head, the torus showed a tuft of silvery scales, internally and the clypeus was without scales; on the anterior leg the V tarsus has smooth nails.

In the larva (Figure 3), short and hyaline lateral spines are prominent on the thorax; on the head bifurcated bristle 6 and multiple bristle 7 are visible; in the VIII abdominal segment pecten is in a single row, presenting as a long spine with a small serrated base or fringes on the lateral bases; in the siphon, the pecten is regularly spaced, and the 1S bristle is composed of two to four branches. In the pupa, the swimming reed possesses long hairs along its edge; in the abdomen, seta 1 shows dichotomous ramifications; bristle 9 is simple, with small side hairs. All these identification features of the morphology of *Ae. albopictus*, adhered to the classification of the ^6^Ministry of Health (1989) and ^7^Consoli & Lourenço-de-Oliveira (1994).

The specimens of one larva and one pupa per slide, prepared on two separate slides, plus the four adult females were deposited in the Entomological Reference Collection of the Department of Epidemiology, Faculty of Public Health, University of São Paulo.

The presence of *Ae. albopictus* in Acre reinforces the rapid expansion of this species in Brazil. In the Americas, including Brazil, *Ae. albopictus* is considered a potential vector of the dengue arboviruses (DENV), Zika (ZIKV), chikungunya (CHIKV) and yellow fever virus (YFV)^8^. Besides, the presence of vertical transmission of these arboviruses in the *Ae. albopictus* females has already been reported under natural conditions, both in Brazil and worldwide^5,8,9,10^. Vertical transmission appears to be an important mechanism for maintaining the circulation of the arboviruses during the less favorable periods of transmission. The occurrence of this phenomenon in the *Ae. albopictus* populations in countries where it is not regarded as a primary vector of arboviruses, such as Brazil, raises an alert for the possible role that this species may play in the epidemiological scenario of the transmission of these pathogens^5,9,10^.

In Brazil, *Ae. albopictus* is present more abundantly in the sylvatic and rural areas, although its presence has already been recorded in the suburban and urban areas, with immature forms occurring both in the natural and artificial breeding sites^5.11,12^. In parallel, *Ae. albopictus* presents eclecticism in relation to the choice of which vertebrate host to blood-feed upon, feeding on both human beings, as well as other vertebrate animals^5,8^. Such behavior may favor this species as a bridge vector for the arboviruses that circulate in the sylvatic areas, such as YFV^13^.

A recent study using secondary data from entomological surveillance in Brazil indicated the presence of *Ae. albopictus* in the state of Acre, since 2020^14^, but without an official record of any findings of the immature forms or adults of this species. The methods used for the surveillance of *Aedes* with medical importance, especially *Aedes aegypti*, are highly dependent on human operation and skills to identify the mosquito species, both in the field and laboratory; therefore, it is important to invest time and resources in the training and qualification of health agents^15^.

This paper shows, for the first time, that *Ae. albopictus* is present in all the Brazilian states, including Acre. With the record of *Ae. albopictus* in Rio Branco, it is interesting to investigate the presence of this species in the other municipalities in Acre. Besides, the presence of this species encourages new studies on the expansion and bioecology of this vector, in the state.

In the current context, it is important to include identification of the species *Ae. Albopictus*, in the routine activities performed by the endemic agents. The maintenance of continuous entomological and virological surveillance of the *Ae. albopictus*, now recorded in all the Brazilian states, is essential to detect any possible change in the role of this species in the dynamics of the transmission of DENV, ZIKV, CHIKV and YFV, in Brazil.

## Acknowledgments

The authors thank to the Rio Branco City Hall for logistical support.

